# Choice of host model and manipulated transcription regulator dictates T6SS effector-mediated toxicity

**DOI:** 10.64898/2026.02.11.705042

**Authors:** Hadar Cohen, Eyal Elias, Dan Bar Yaacov, Dor Salomon, Motti Gerlic

**Affiliations:** Gray School of Medical Sciences, Gray Faculty of Medical and Health Sciences, Tel Aviv University, Tel Aviv, Israel; The Shraga Segal Department of Microbiology, Immunology, and Genetics, Ben-Gurion University of the Negev, Beer-Sheva, Israel

## Abstract

Type VI secretion system (T6SS), a toxic effector-delivery apparatus primarily studied for its antibacterial properties, has recently emerged as a widespread anti-eukaryotic determinant, particularly in members of the genus *Vibrio*. Although various anti-eukaryotic effectors have been described, it remains unknown whether effectors target distinct hosts in a similar manner. Moreover, it is unclear whether the entire effector arsenal encoded within a single genome is co-regulated under different conditions and by different regulators. Here, we employ the anti-eukaryotic T6SS3 of *Vibrio proteolyticus* to address these knowledge gaps. By monitoring toxicity during infection of oyster hemocytes and comparing the results with previous infections of murine macrophages, we find that the effector Tie1 plays a role only in intoxicating murine cells, whereas Tie2 intoxicates both cell models. Furthermore, we show that artificial induction of T6SS3 via deletion of the negative regulator *hns1* fails to induce the orphan effector Tie3, whereas overexpression of a T6SS3-specific activator, Ats3, induces all three known effectors. Ats3 overexpression further reveals that Tie3, not only Tie2, participates in hemocyte intoxication. Collectively, these findings indicate that analyses of T6SS-mediated infections must factor in the possibility of partial effector repertoire activation even when the main gene cluster is fully induced, and that the contribution of anti-eukaryotic T6SS effectors to intoxication is host-dependent.

## Main Text

Many Gram-negative bacteria deploy the Type VI secretion system (T6SS), a toxic effector delivery apparatus, as an antibacterial tool to mediate interbacterial competition [1, 2]. However, recent work revealed a previously underappreciated role for T6SSs in interactions with eukaryotes, particularly among marine bacteria belonging to the genus *Vibrio* [3–5]. One such example is T6SS3 in *Vibrio proteolyticus* (*Vpr*), which, as we have shown, is used to intoxicate the aquatic model organism *Artemia salina* [6] and to induce inflammasome-dependent cytotoxicity in a mammalian model immune cell, murine bone marrow-derived macrophages (BMDMs) [7]. *Vpr* T6SS3 secretes at least three anti-eukaryotic effectors: Tie1, Tie2, and Tie3 (ACWZF4_RS09470, ACWZF4_RS09465, and ACWZF4_RS22085, respectively). We previously established that Tie1 and Tie2, both encoded within the T6SS3 gene cluster, drive pyroptotic cell death in BMDMs [7], while the orphan effector Tie3 works in concert with Tie1 and Tie2 to induce *Artemia* nauplii lethality [6].

In previous studies, we drew inspiration from earlier T6SS works [8–11] and manipulated positive (i.e., the T6SS3 regulator Ats3) or negative (i.e., the histone-like nucleoid-structuring repressor H-NS1) transcription regulators to induce T6SS3 expression levels and enhance T6SS-mediated toxic phenotypes [7]. However, it remains unknown whether manipulating different regulators can differentially affect T6SS-mediated toxicity and whether T6SS effectors have similar effects across distinct host models. To address these knowledge gaps, we set out to compare the effect of *Vpr* T6SS3 and the different regulators during infection of immune cells from a more habitat-relevant marine organism (i.e., hemocytes), the oyster *Crassostrea gigas*, with our previous observations using the BMDM model [7].

First, we used real-time microscopy (Incucyte) to monitor the toxicity of *Vpr* strains in which we induced T6SS3 by deleting the negative regulator *hns1* (ACWZF4_RS09750) during hemocyte infection. As we observed in BMDM infection [7], a Δ*hns1* strain induced dramatic hemocyte cell death that was completely dependent on T6SS3, as an isogenic T6SS3^−^ strain induced no cell death (Figure 1A-B). Interestingly, the deletion of the effector *tie2*, alone or in combination with other effectors, was sufficient to abrogate toxicity in hemocytes. However, in previous work, we showed that both Tie1 and Tie2 contribute to T6SS3-mediated cell death in BMDMs [7]. Taken together, these results demonstrate that Tie1 does not significantly contribute to hemocyte toxicity, thus suggesting that T6SS effectors may contribute differently to infection, depending on the host cell investigated.

**Figure 1.**
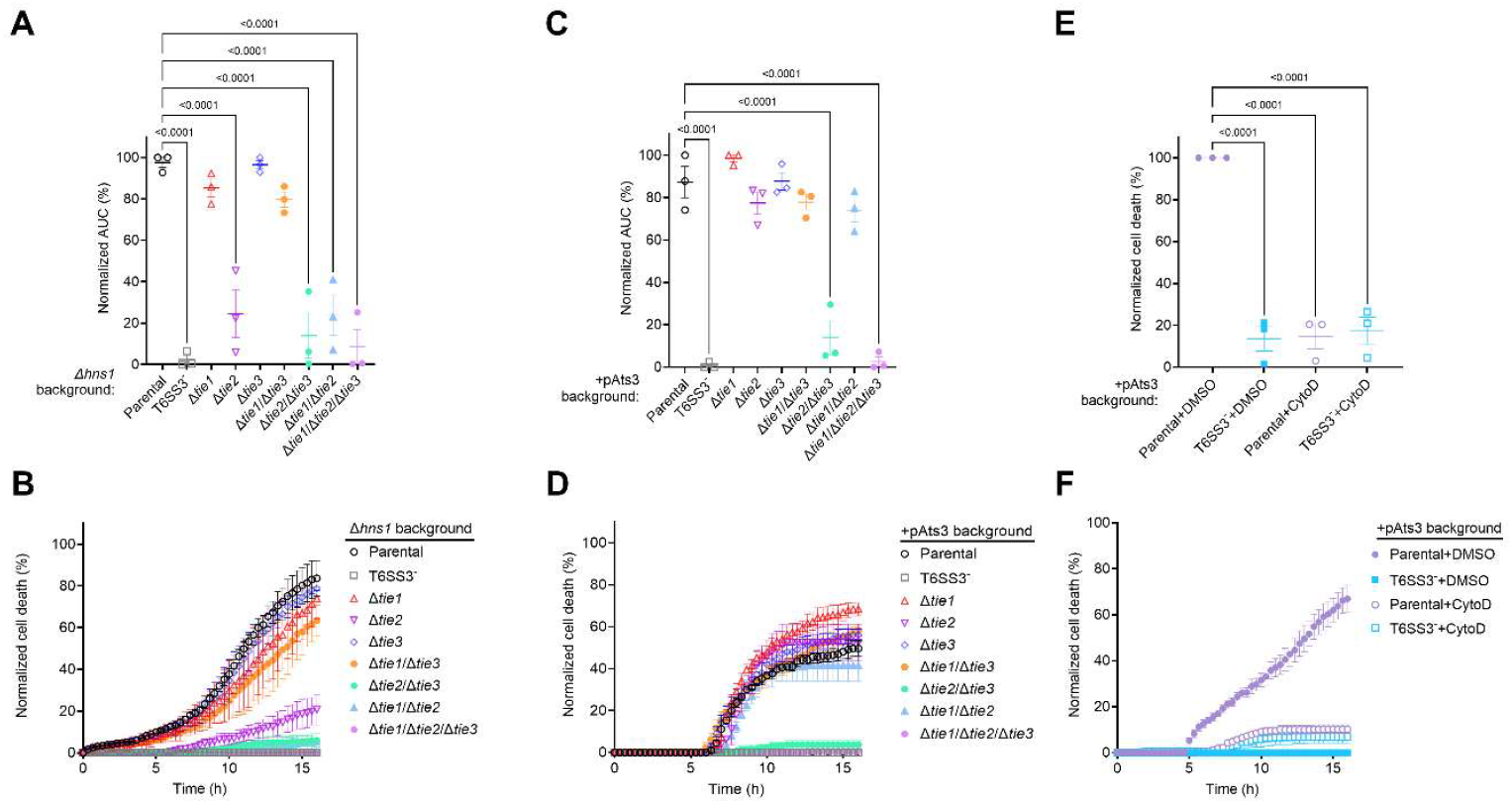
T6SS3 effector contribution to toxicity during oyster hemocyte infection depends on the inducing regulatory pathway. **(A-F)** Approximately 2×10^5^ hemocytes extracted from *C. gigas* hemolymph were seeded into 96-well plates coated with poly-D-lysine in Instant Ocean media in triplicate. Cells were infected with the indicated *Vpr* Δ*vprh* strains at a multiplicity of infection (MOI) = 5. Kanamycin was added to the media to maintain an arabinose-inducible plasmid for the expression of Ats3 (pAts3), and arabinose (0.1% [w/v]) was added to induce expression from the P*bad* promoter (C-F). When indicated (E-F), cytochalasin D (Cyto D; 5 μM) was added to the cells 30 minutes prior to infection; an equivalent volume of DMSO was added as a control. Propidium iodide (PI) uptake over 16 hours was assessed by real-time microscopy (Incucyte S3) as an indicator of cell death, and the data were recorded as PI-positive cells normalized to cell confluency at t = 0. The data (mean ± SD) of three independent experiments were then grouped as the area under the curve (AUC) (A, C, E). The cell death kinetics of representative experiments are shown in B, D, and F (mean ± SE; n = 3 infection wells). Statistical comparisons in A, C, and E between samples at the 16-hour timepoint were performed using a one-way ANOVA followed by Dunnett’s multiple comparison test; significant differences (*P* < 0.05) are denoted only for comparisons against the parental strain.

To further dissect the virulence activity of T6SS3 by engaging a different regulatory pathway, we infected oyster hemocytes with *Vpr* strains overexpressing Ats3. Real-time microscopy revealed that, as with *hns1* deletion, Ats3-overexpression induced a T6SS3-dependent cell death (Figure 1C-D). Notably, this T6SS3-induced hemocyte cell death is dependent on bacterial internalization and was abolished in the presence of cytochalasin D, which inhibits phagocytosis by blocking actin polymerization [12, 13], as previously shown in BMDMs [7] (Figure 1E-F). Surprisingly, in contrast to the pivotal contribution of Tie2 to hemocyte toxicity in strains in which T6SS3 is induced by *hns1* deletion, Ats3-induced T6SS3-dependent hemocyte cell death was not completely dependent on Tie2. Instead, the orphan effector Tie3 was sufficient to induce hemocyte cell death in the absence of Tie2, and only the deletion of both *tie2* and *tie3* abolished the T6SS3-dependent cell death (Figure 1C-D). Together, these results imply that *hns1* deletion and Ats3 overexpression lead to hemocyte cell death that depends on distinct effector subsets. This observation prompted the hypothesis that induction of T6SS3 by distinct transcriptional regulators results in distinct effector repertoires.

To test our hypothesis, we performed RNA-seq analyses and compared the effect of *hns1* deletion and Ats3 overexpression on the *Vpr* transcriptional landscape (Supplementary Dataset S1). Consistent with our hypothesis, the overexpression of Ats3 induced the expression of all known T6SS3 effectors, including the orphan effector *tie3*, whereas the deletion of *hns1* failed to induce *tie3* expression (Figure 2A-C). Thus, this difference in *tie3* expression accounts for the observed difference in its contribution to host toxicity between the *hns1* deletion and Ats3 overexpression strains. Although manipulation of both regulators upregulated the genes comprising the T6SS3 main cluster, Ats3 overexpression yielded a greater fold change than *hns1* deletion (Figure 2C). Furthermore, *hns1* deletion resulted in a broad transcriptional reprogramming, affecting multiple pathways, including the upregulation of the antibacterial T6SS1 (Figure 2B, D and Supplementary Dataset S2); in contrast, the Ats3 regulon was more specific to T6SS3 upregulation (Figure 2A, D and Supplementary Dataset S3). Collectively, these results reveal distinct upregulation of T6SS3 effectors by different transcriptional cascades, thereby explaining the differential reliance on specific effectors during host cell infection.

**Figure 2.**
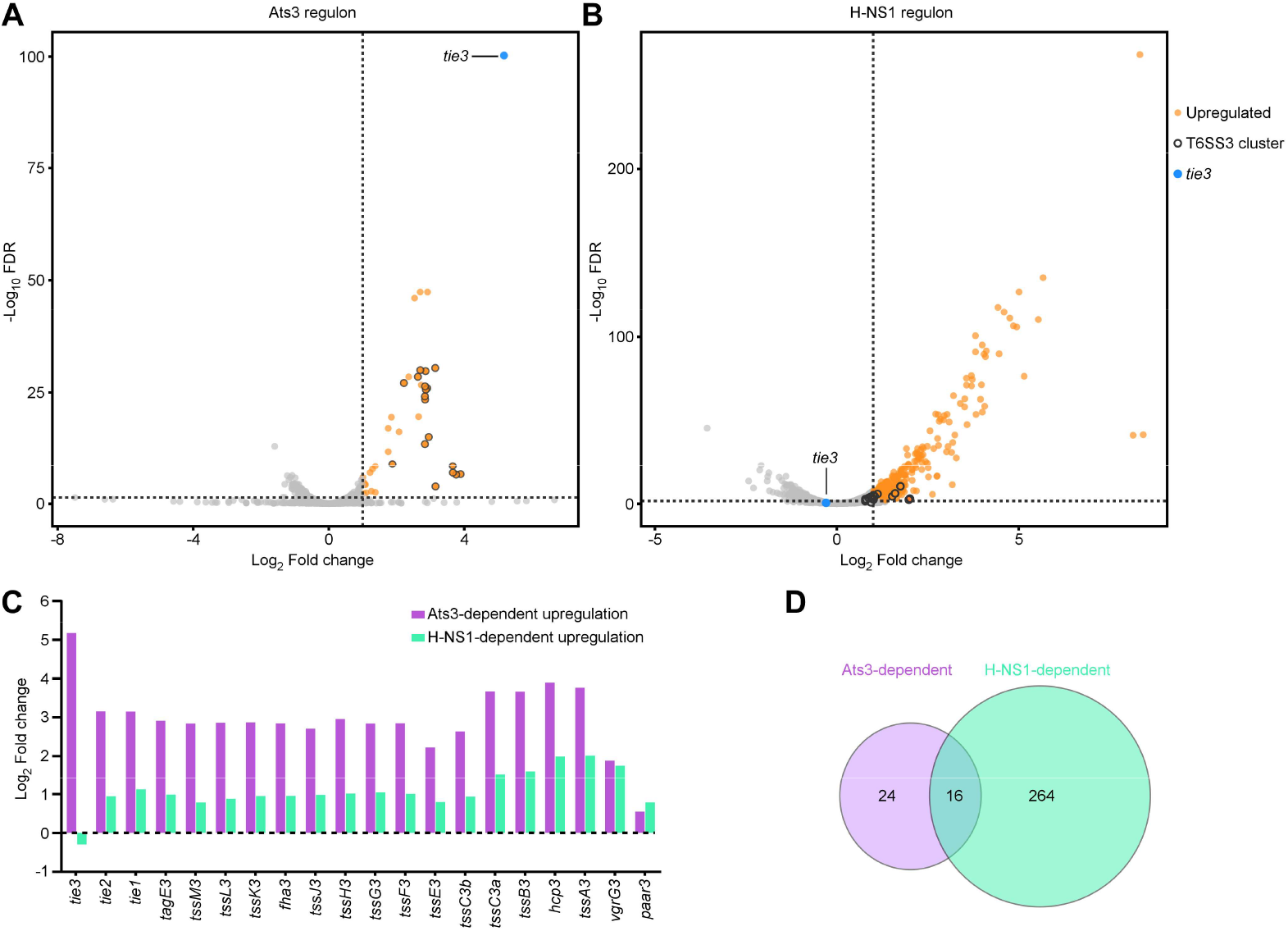
Ats3 overexpression, but not *hns1* deletion, upregulates *tie3* transcripts. **(A–B)** Volcano plots of the RNA-seq data are shown for wild-type *Vpr* overexpressing Ats3 from a plasmid (pAts3; A) or the Δ*hns1* derivative carrying an empty plasmid (pEmpty; B), each compared to wild-type *Vpr* carrying pEmpty. The artificially manipulated genes (*ats3* in A and *hns1* in B) were excluded from the plots for clarity. Significantly upregulated genes (orange) were considered as having a fold change > 2 and a false discovery rate (FDR) < 1%. (**C**) Comparison of T6SS3 gene expression changes in *Vpr* containing pAts3 or in a Δ*hns1* derivative containing pEmpty; fold change was calculated relative to *Vpr* containing pEmpty. (**D**) Venn diagram showing the number of genes significantly upregulated (fold change ≥ 2; FDR ≤ 0.01) by Ats3 overexpression and *hns1* deletion.

## Conclusions

In this work, we elucidate the role of *Vpr* T6SS3 during infection of immune cells from a marine animal model. Although we previously showed that the effectors Tie1 and Tie2 promote BMDM cell death, here we demonstrate that the effectors Tie2 and Tie3, but not Tie1, are key mediators of hemocyte cell death. These differences highlight the importance of selecting an appropriate host model when assessing the contributions of effector proteins to virulence. Notably, we cannot rule out the possibility that different media and temperature conditions, rather than different host cells, contributed to the effect of Tie1 during infection of BMDMs and oyster hemocytes. Furthermore, although *hns1* deletion upregulated T6SS3 activity and expression of effectors within the main cluster, it did not induce the expression of the orphan effector *tie3*. In contrast, activation via Ats3, a T6SS3-specific regulator, upregulated *tie3* expression. These findings emphasize the critical role of the activation mode in shaping the functional T6SS effector repertoire, particularly in systems that harbor orphan effectors and when the T6SS is induced by artificial regulator manipulations.

## Supporting information

Supplementary Materials and Methods, Tables, Dataset captions, and References

## Data availability

The *Vpr* genome assembly was deposited to NCBI under BioProject PRJNA1373524 (assembly GCA_053890595.1). RNA-seq data generated in this study were deposited to the NCBI SRA under accession PRJNA1422434.

## Acknowledgements

This project received funding from the US-Israel Binational Science Foundation (https://www.bsf.org.il; award number 2021733 to DS), from the European Research Council (Project 101116636 — REDBAC to DB), and from the Israel Science Foundation (www.isf.org.il; ISF grant number 2174/22 to MG).

## Author Contributions

Conceptualization, D.S., M.G., and H.C.; Methodology, H.C, and E.E.; Investigation, H.C., E.E., and D.B.; Formal analysis, H.C., D.B., D.S., and M.G.; Writing – Original Draft, H.C.; Writing – Review & Editing, D.B., D.S., and M.G.; Funding acquisition, D.B., D.S., and M.G.; Supervision, D.S. and M.G.

