## Supplementary Materials and Methods, Tables, Dataset captions, and References for "Choice of host model and manipulated transcription regulator dictates T6SS effector-mediated toxicity"

**Supplementary Tables S1-S2**

**Supplementary Datasets S1-S3 (captions)**

**Supplementary References**

### Supplementary Materials and Methods

#### Bacterial strains and media

For a complete list of strains and plasmids used in this study, see Supplementary Tables S1 and S2, respectively. *V. proteolyticus* ATCC 15338 (*Vpr*) and its derivative strains were grown in MLB (Lysogeny broth supplemented with NaCl to a final concentration of 3% [w/v]) or on MLB agar (1.5% [w/v]) plates at 30°C. When plasmid maintenance was required, media were supplemented with kanamycin (250 µg/mL). To induce the expression from pBAD plasmids, 0.1% (w/v) L-arabinose was added to the media.

#### Construction of bacterial deletion strains

*E. coli DH5α* (λ-pir) strains carrying pDM4 [1] suicide plasmids for gene deletions were used as donor strains to introduce the plasmid into *Vpr* via conjugation. Trans-conjugants were selected on MLB agar plates containing chloramphenicol (10 µg/mL). The resulting trans-conjugants were then grown on MLB agar plates containing sucrose (15% [w/v]) for counter-selection and loss of the *sacB*-containing pDM4. Deletions were confirmed by PCR.

#### Construction of bacterial overexpression strains

*E. coli DH5α* (λ-pir) carrying pBAD/Myc-His$^{Kan}$ constructs were used as donor strains to introduce the plasmids into *Vpr* via conjugation. Trans-conjugants were selected and maintained on MLB agar plates supplemented with kanamycin (250 µg/mL).

#### Hemocyte collection and processing

*Crassostrea gigas* (*C. gigas*) were obtained from a local supplier (Batshon Fish) one day prior to hemolymph collection. Hemolymph was harvested as previously described [2], with minor modifications. Briefly, oyster shells were opened carefully with an oyster knife. Using a cold 1 mL syringe fitted with a 25G needle, hemolymph was withdrawn from the adductor muscle, taking care to avoid seawater contamination, and immediately transferred into a cold Falcon tube on ice. Each sample was examined under a light microscope to assess quality; only samples with no visible debris and minimal hemocyte aggregation were used and combined into a single sample.

Cell samples were centrifuged at 400 × *g* for 10 minutes at 4°C, and the supernatants were discarded. Cells were resuspended in Instant Ocean media (IO; 3.3% [w/v]; Aquarium Systems) and counted under a microscope. Approximately $2 \times 10^5$ cells were seeded per well in 96-well plates coated with poly-D-lysine (Nu-152039, ThermoFisher). The seeded cells were centrifuged at 400 × *g* for 5 minutes at 4°C, and then incubated at room temperature for at least one hour to allow adherence before bacterial infection.

**Infection assays**

Hemocyte media were replaced with fresh IO supplemented with propidium iodide (PI; 1 µg/mL; Sigma-Aldrich #P4170). For phagocytosis inhibition assays, cytochalasin D (final concentration 5 µM; Biotest #1233) was added 30 minutes prior to infection. When plasmid maintenance and arabinose-induced expression were required, kanamycin (250 µg/mL) and L-arabinose (0.1% [w/v]) were added to the media.

Hemocytes were infected with *Vpr* strains at a multiplicity of infection (MOI) of 5. To this end, overnight cultures of *Vpr* were diluted two-fold and, when required, kanamycin and L-arabinose were added to the media. After three hours of growth, bacterial cultures were normalized to $OD_{600}$ = 0.016 in IO. Then, 50 µL of the normalized bacterial cultures were added to wells containing the hemocytes, and plates were centrifuged for 5 minutes at 400 × *g* to initiate infection. The plates were then placed in Incucyte SX5 (Sartorius) and incubated at 28°C; images were acquired every 20 minutes, and cell death kinetics were analyzed using the Incucyte 2024B analysis software and then exported to the GraphPad Prism software. Normalization was performed according to the maximal PI-positive object count to calculate the percentage of dead cells [3].

**Genomic DNA Sequencing**

Genomic DNA (gDNA) was extracted from *Vpr* ATCC 15338 using the Presto Mini gDNA Bacteria Kit (Geneaid), following the manufacturer's instructions. The gDNA was sequenced with Oxford Nanopore Technology using long-read sequencing technology by Plasmidsaurus (https://www.plasmidsaurus.com/), including construction of an amplification-free long-read sequencing library using v14 library prep chemistry followed by library sequencing using a primer-free protocol [4]. The genome assembly was deposited to NCBI under BioProject PRJNA1373524 (assembly GCA_053890595.1) and annotated using the NCBI Prokaryotic Genome Annotation Pipeline (PGAP).

**Differential gene expression analysis**

*Bacterial culture preparation:* *Vpr* strains were grown in triplicate in MLB media, with each biological replicate initiated on a different day from a single colony. At mid-log phase ($OD_{600}$ = 4.5), 1 mL of culture was collected, the cells were harvested by centrifugation, and the cell pellets were flash-frozen and kept at −80°C.

*RNA extraction:* The supernatants were discarded, and RNA was extracted from the bacterial pellets using the GeneJET RNA Purification Kit (Thermo Scientific #K0731). RNA samples were treated with 4 units of DNase I (NEB # M0303L) for 20 minutes at 37°C. RNA integrity was determined on a QIAxcel device (QIAGEN, QIAxcel RNA QC Kit v2.0), and concentration was determined using a QuantiFluor® RNA System (Promega, #E3310) on a Qubit™ Flex Fluorometer (Invitrogen).

*RNA-seq analyses:* RNA-seq libraries were constructed using the Zymo-Seq RiboFree Total RNA Library Kit per manufacturer's instructions (Zymo Research, #R3003). The molarity of libraries was determined using QIAxcel (QIAxcel DNA High Sensitivity Kit) and QuantiFluor® dsDNA System (Promega, # E2670) on a Qubit™ Flex Fluorometer (Invitrogen). Libraries were sequenced (150PE) on a NovaSeq X (Illumina).

Genomics Workbench was used for all analysis steps as described below. RNA-seq reads were first trimmed according to length and quality scores to ensure high quality of the reads by using the following parameters: Trim using quality scores = Yes; Quality limit = 0.01; Trim ambiguous nucleotides = Yes; Maximum number of ambiguities = 1; Automatic read-through adapter trimming = Yes; Trim homopolymers from 5' = No; Trim homopolymers from 3' = Yes; polyA = No; polyC = No; polyG = Yes; polyT = No; Remove 5' terminal nucleotides = No; Remove 3' terminal nucleotides = No; Remove on first read = Yes; Remove on second read (for paired reads) = Yes; Trim to a fixed length = No; Maximum length = 150; Trim end = Trim from 3'-end; Discard short reads = Yes; Minimum length = 50; Discard long reads = No; Save discarded sequences = No; Save broken pairs = No.

Next, we performed differential gene expression analysis. To obtain normalized expression measurements (as transcripts per million, TPM), we mapped the RNA-seq reads (following quality control and trimming of adapter read-through) to the Genome of *Vibrio proteolyticus* ATCC 15338 (NCBI RefSeq NZ_CM132682.1 and NZ_CM132683.1) with the following parameters in CLC Genomics Workbench: Reference type = Genome annotated with genes only; Reference sequence = *Vibrio proteolyticus* strain ATCC 15338; Gene track = *Vibrio proteolyticus* strain ATCC 15338_Gene; Use spike-in controls = no; Mismatch cost = 2; Insertion cost = 3; Deletion cost = 3; Length fraction = 0.8; Similarity fraction = 0.8; Global alignment = No; Strand specific = Reverse; Library type = Bulk; Maximum number of hits for a read = 10; Count paired reads as two = No; Ignore broken pairs = No; Expression value = TPM. Finally, we analyzed differential gene expressions using default parameters (also using CLC Genomics Workbench).

**Statistical analyses**

The data were analyzed using GraphPad Prism 9. Data are presented as the mean ± SE or SD, as indicated. Comparisons were performed using RM one-way ANOVA, followed by Dunnet's multiple comparison, unless otherwise stated. Statistical significance was considered at $P < 0.05$.

### Supplementary Tables

##### Supplementary Table S1: A list of bacterial strains used in this study

| Strain name | Genotype | Source | Comments |
|---|---|---|---|
| *Vibrio proteolyticus* ATCC 15338 | Wild-type | ATCC collection | Used for differential gene expression analysis |
| Δ*vprh* | *Vibrio proteolyticus* ATCC 15338 Δ*vprh* | [5] | Used for hemocyte infection |
| Δ*vprh*/T6SS3$^{-}$ | *Vibrio proteolyticus* ATCC 15338 Δ*vprh*/Δ*tssL3* | [6] | Used for hemocyte infection |
| Δ*vprh*/Δ*tie1* | *Vibrio proteolyticus* ATCC 15338 Δ*vprh*/Δ*tie1* | [7] | Used for hemocyte infection |
| Δ*vprh*/Δ*tie2* | *Vibrio proteolyticus* ATCC 15338 Δ*vprh*/Δ*tie2* | [7] | Used for hemocyte infection |
| Δ*vprh*/ Δ*tie3* | *Vibrio proteolyticus* ATCC 15338 Δ*vprh*/Δ*tie3* | [7] | Used for hemocyte infection |
| Δ*vprh*/Δ*tie1*/Δ*tie2*/Δ*tie3* | *Vibrio proteolyticus* ATCC 15338 Δ*vprh*/Δ*tie1*/Δ*tie2*/Δ*tie3* | [7] | Used for hemocyte infection |
| Δ*vprh*/Δ*tie1*/Δ*tie2* | *Vibrio proteolyticus* ATCC 15338 Δ*vprh*/Δ*tie1*/Δ*tie2* | This study | Used for hemocyte infection |

| | | | |
|---|---|---|---|
| Δ*vprh*/Δ*tie1*/Δ*tie3* | *Vibrio proteolyticus* ATCC 15338 Δ*vprh*/Δ*tie1*/Δ*tie3* | This study | Used for hemocyte infection |
| Δ*vprh*/Δ*tie2*/Δ*tie3* | *Vibrio proteolyticus* ATCC 15338 Δ*vprh*/Δ*tie2*/Δ*tie3* | This study | Used for hemocyte infection |
| Δ*vprh*/Δ*hns1* | *Vibrio proteolyticus* ATCC 15338 Δ*vprh*/ Δ*hns1* | [6] | Used for hemocyte infection |
| Δ*vprh*/Δ*hns1*/T6SS3$^{-}$ | *Vibrio proteolyticus* ATCC 15338 Δ*vprh*/Δ*hns1*/Δ*tssL3* | [6] | Used for hemocyte infection |
| Δ*vprh*/Δ*hns1*/Δ*tie1* | *Vibrio proteolyticus* ATCC 15338 Δ*vprh*/Δ*hns1*/Δ*tie1* | [6] | Used for hemocyte infection |
| Δ*vprh*/Δ*hns1*/Δ*tie2* | *Vibrio proteolyticus* ATCC 15338 Δ*vprh*/Δ*hns1*/Δ*tie2* | [6] | Used for hemocyte infection |
| Δ*vprh*/Δ*hns1*/Δ*tie1*/Δ*tie2* | *Vibrio proteolyticus* ATCC 15338 Δ*vprh*/Δ*hns1*/Δ*tie1*/Δ*tie2* | [6] | Used for hemocyte infection |
| Δ*vprh*/Δ*hns1*/Δ*tie3* | *Vibrio proteolyticus* ATCC 15338 Δ*vprh*/Δ*hns1*/Δ*tie3* | [7] | Used for hemocyte infection |
| Δ*vprh*/Δ*hns1*/Δ*tie1*/Δ*tie2*/ Δ*tie3* | *Vibrio proteolyticus* ATCC 15338 Δ*vprh*/Δ*hns1*/Δ*tie1*/Δ*tie2*/Δ*tie3* | [7] | Used for hemocyte infection |
| Δ*vprh*/Δ*hns1*/Δ*tie1*/Δ*tie3* | *Vibrio proteolyticus* ATCC 15338 Δ*vprh*/Δ*hns1*/Δ*tie1*/Δ*tie3* | This study | Used for hemocyte infection |

| | | | |
|---|---|---|---|
| Δ*vprh*/Δ*hns1*/Δ*tie2*/Δ*tie3* | *Vibrio proteolyticus* ATCC 15338 Δ*vprh*/Δ*hns1*/Δ*tie2*/Δ*tie3* | This study | Used for hemocyte infection |
| Δ*hns1* | *Vibrio proteolyticus* ATCC 15338 Δ*hns1* | This study | Used for differential gene expression analysis |
| *Escherichia coli* DH5α (λ-pir) | K-12 derivative laboratory strain containing λ pir | Obtained from Eric V. Stabb | Used for plasmid construction and maintenance |
| *Escherichia coli* HB101 | K-12 derivative laboratory strain | ATCC collection | Harbors pRK2013; used as a conjugation helper strain |

**Supplementary Table S2. A list of plasmids used in this study**

| Plasmid name | Description | Comments | Source |
|---|---|---|---|
| pBAD/Myc-His$^{Kan}$ | pBR322 ori-containing plasmid harboring a Kan$^{R}$ cassette, araC, and an MCS following a P*bad* promoter | Used as an empty control plasmid | [8] |
| pAts3 | pBAD/Myc-His$^{Kan}$ containing the *ats3* ORF in its MCS, not fused to the C-terminal tag encoded on the plasmid | Used for arabinose-inducible expression of Ats3 | [6] |
| pDM4:*tie3* | pDM4 suicide plasmid containing 1 kb upstream and 1 kb downstream of *tie3* in its MCS | Used to delete *tie3* in *V. proteolyticus* | [7] |
| pDM4:*hns1* | pDM4 suicide plasmid containing 1 kb upstream and 1 kb downstream of *hns1* in its MCS | Used to delete *hns1* in *V. proteolyticus* | [6] |

#### Supplementary Datasets (captions)

**Supplementary Dataset S1. RNA-seq results**

**Supplementary Dataset S2. Transcripts significantly upregulated upon *hns1* deletion**

**Supplementary Dataset S3. Transcripts significantly upregulated upon *ats3* overexpression**
